# Quantifying anti-DUX4 therapy for facioscapulohumeral muscular dystrophy

**DOI:** 10.1101/2024.08.14.607973

**Authors:** Matthew V. Cowley, Peter S. Zammit, Christopher R. S. Banerji

## Abstract

Facioscapulohumeral muscular dystrophy (FSHD) is an inherited skeletal myopathy with no cure. Expression of the myotoxic transcription factor double homeobox 4 (*DUX4*) is believed to underlie FSHD pathogenesis and many proposed therapies target DUX4 generation or function. Which of these therapies will be the most effective is unclear. Here, by constructing a Markov-chain-based mathematical model of DUX4-mediated myotoxity in FSHD, we interrogate various anti-DUX4 FSHD therapeutic strategies. We derive an analytical function for myonuclear life expectancy in terms of the parameters of *DUX4* expression and function. In a biologically relevant parameter regime, therapeutically decreasing the DUX4 protein diffusion rate is, surprisingly, predicted to be more effective at increasing myonuclear life expectancy than reducing the rate of myonuclear apoptosis caused by the expression of DUX4-target genes. We find that targeting elements of *DUX4* transcription/translation, such as mRNA stability via siRNA therapy, has a limited predicted impact on DUX4-meditated toxicity when performed in isolation. However, our model predicts a super-additive effect from combining transcription/translation targeting strategies with approaches that minimise DUX4 diffusion-mediated import into neighbouring myonuclei. Importantly, we provide a computational tool to test and inform therapeutic designs, enabling pre-clinical screening of FSHD treatment approaches.

## I. INTRODUCTION

Facioscapulohumeral muscular dystrophy (FSHD) is a progressive and debilitating, inherited myopathy that has no cure [1, 2]. FSHD is autosomal dominant, causing muscle atrophy and fatty tissue replacement of skeletal muscle [3]. As the condition progresses, between 8 and 23 % of patients will become wheelchair dependant, motivating studies to accelerate progress towards treatment [4–6]. The transcription factor double homeobox 4 (DUX4) is strongly linked to FSHD pathology [7, 8], and many proposed therapies target elements of the *DUX4* pathway. These can be broadly grouped into strategies that target *DUX4* transcription, *DUX4* mRNA stability, *DUX4* translation, DUX4 protein stability, or DUX4 target gene activation [9].

*DUX4* transcription inhibitors include: losmapimod, a p38*α* inhibitor currently in phase 3 clinical trials (NCT05397470) [10], and molecules targeting regulation upstream of the *DUX4* promoter [11]. Approaches targeting *DUX4* transcript stability include various RNA interference strategies such as the antibody-oligonucleotide conjugate AOC 1020 currently in phase 1/2 clinical trials (NCT05747924) [12], siRNAs [13], and LNA gapmers [14]. DUX4 target gene activation is inhibited via approaches such as aptamers that can bind DUX4 [15], or p300 inhibitors which suppress DUX4-mediated chromatin availability [16]. The comparative potential efficacy of these diverse routes has yet to be objectively evaluated.

We recently proposed an *in silico* model for the role of *DUX4* in FSHD, describing the condition through a compartment model: a set of deterministic ordinary differential equations (ODEs) parameterised from experimental data [17]. These equations describe the progression of a myocyte from healthy states, with no *DUX4* transcripts or downstream target genes of DUX4 (D4Ts) present, to those with transcriptional activity of *DUX4* and D4Ts, and finally to apoptotic states (Figure 1**A**). The transitions between these states are described by the parameters outlined in Table I, and are calculated from experimental data. The transcription rates are calculated by fitting a two-step promotor model to single-cell mRNA counts, the degradation rate from fitting an exponential decay to RT-qPCR time series data, the translation rate from delay time in RT-qPCR experiments, the apoptosis rate from a GFP reporter assay, and the syncytial import rate from fitting cell phenotypes in single-cell data to a cellular automaton of an FSHD myofibre. For further details please see our previous work [17].

**Figure 1.**
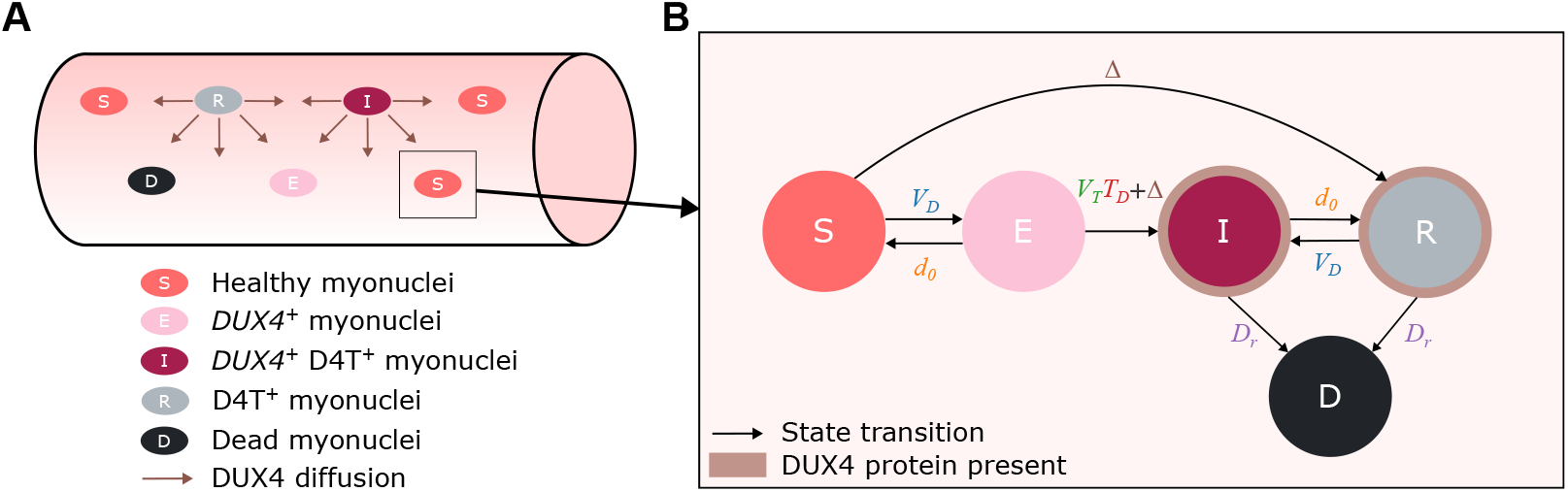
The *DUX4* pathway in facioscapulohumeral muscular dystrophy can be modelled via a continuous time Markov chain of a myonucleus in an affected myofibre. (**A**) Schematic showing a myofibre with brown arrows indicating protein transport between myonuclei. States are assigned via transcriptional signature and D4T-positivity necessitates DUX4 protein-positivity [17]. Therefore all D4T-positive myonuclei can act as a source of DUX4. **(B)** Transition diagram of a continuous-time Markov chain of a single myonucleus participating in the *DUX4* pathway with four transcriptomic states (susceptible *S* (*DUX4* ^−^ D4T^−^), exposed *E* (*DUX4* ^+^ D4T^−^), infected *I* (*DUX4* ^+^ D4T^+^), resigned *R* (*DUX4* ^−^ D4T^+^)) and and one dead state *D*. Six rate parameters govern the transitions between the states as shown by the black arrows. States *I* and *R* have DUX4 protein present as indicated by the brown outline. D4T, DUX4 target gene; *V*_*D*_, *DUX4* transcription rate; *d*_0_, *DUX4* mRNA degradation rate; *V*_*T*_, D4T transcription rate; *T*_*D*_ *DUX4* mRNA translation rate; *D*_*r*_, D4T^+^ myonuclear apoptosis rate; Δ, DUX4 syncytial import rate.

**Table I.** Experimentally derived rate parameters [17] and their definitions. *θ*_*ε*_, endogenous rate; D4T, DUX4 target gene.

| Parameter | Definition | Value $\theta_\epsilon$ |
| --- | --- | --- |
| $V_D$ | $DUX4$ transcription rate | $2.11 \times 10^{-3} \text{ hour}^{-1}$ |
| $d_0$ | $DUX4$ mRNA degradation rate | $2.46 \times 10^{-1} \text{ hour}^{-1}$ |
| $V_T$ | D4T transcription rate | $6.41 \text{ hour}^{-1}$ |
| $T_D$ | $DUX4$ mRNA translation rate | $7.69 \times 10^{-2} \text{ hour}^{-1}$ |
| $D_r$ | D4T$^+$ myonuclear apoptosis rate | $4.95 \times 10^{-2} \text{ hour}^{-1}$ |
| $\Delta$ | $DUX4$ syncytial import rate | $4.02 \times 10^{-2} \text{ hour}^{-1}$ |

Framing the system into a mathematical model is useful, as each of the parameters represents a biological process that therapeutics may target. The parameter *V*_*D*_ may be effectively reduced via small molecules such as losmapimod, which has been shown to alter the expression of upstream regulators of *DUX4* [10, 18]. Alternatively, antibody-oligonucleotide conjugates are designed to degrade *DUX4* mRNA and therefore will increase the *DUX4* mRNA degradation rate (*d*_0_) [12]. If our *in silico* model is reframed to consider a diseased myofibre instead of individual myocytes, the syncytial multinuclear entity can be modelled by using the same ODEs to represent the progression of myonuclei through an equivalent series of states (Figure 1**A**). In this domain, additional transitions are possible due to a background concentration of DUX4 protein undergoing transport between myonuclei, followed by import into the nucleus (as represented by arrows in Figure 1**A**).

In our previous work [17], coupled ODEs provided a useful, yet analytically complex perspective on the *DUX4* pathway. To make more precise statements about the comparative potential efficacy of treatment strategies, a natural extension is to move to a probabilistic formulation. This statistical paradigm better represents the nature of the biology and provides established analytical tools. Markov chains, a mathematical formalism used for describing stochastic processes [19, 20], have been used in mathematical biology to successfully model the evolution of karyotypes to understand chromosomal instability in tumours [21], to predict the progression of carcinomas and melanomas [22], to detect cellular ageing in *Escherichia Coli* [23], and most relevantly to our work, describe stochastic gene expression [24, 25].

Absorbing Markov chains are a subset of this modelling framework where some states are defined as absorbing. Once entered, the absorbing state cannot be left. They have been used in an ecological setting to predict species extinction [26]: a problem which predicting DUX4-mediated myonucleus population extinction echoes. Absorbing Markov chains are useful in our case as they naturally capture myonuclear apoptosis as an absorbing state and have many established analytical properties, allowing us to compute the expected time to myonuclear apoptosis (absorption) precisely, in terms of the parameters of our model [27].

Here, we use an absorbing Markov chain formulation of our model of an FSHD muscle fibre [17] to test different anti-DUX4 strategies for FSHD and so suggest which single-target and combination therapies should be prioritised to prolong the life expectancy of muscle cells in FSHD.

## II. APPROACH

We formulated our syncytial compartment model of a population of myonuclei in an individual with FSHD [17] into a continuous-time Markov chain (Figure 1**B**). This Markov chain is constructed of a set of holding times defining the time spent in each state and an embedded discrete-time Markov chain known as the jump chain. As the holding times for each state are unknown, we assume they are fixed and comparable. This permits the treatment of the possible transitions as competing independent exponential random variables [28, 29]. These can be viewed as “exponential clocks”, an approach which helpfully allows the determination of transition probabilities from our experimentally derived state transition rates in Table I [17].

Given that we start in some state *i*, there are *k* possible transitions where *k* is the number of arrows from state *i* in Figure 1**B**. The time taken to transition from state *i* to state *j* is a random variable with an exponential distribution 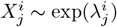, where 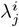 is the rate of the state transition from *i* to *j* in the compartment model. The time to the first state transition from *i* to any other state is simply

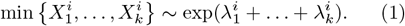

The probability that the first transition from *i* is to a given state *j* can therefore be formulated as

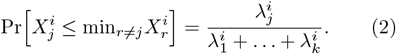

This means that each transition probability is the quotient of the transition rate and the sum of possible transition rates.

The probability matrix of the discrete-time Markov chain is then given by,

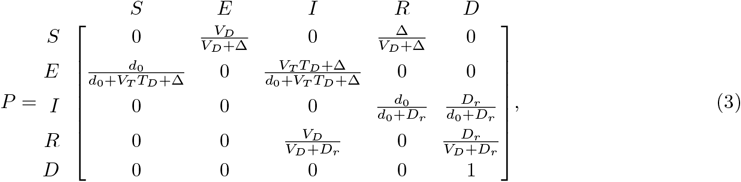

where the rows represent the starting states and the columns represent the first transition states. The (*i, j*)^*th*^ element of *P* is the probability that the first transition from state *i* is to state *j*, with the rates describing the transitions shown in Figure 1**B** and quantified in Table I.

The states in *P* are susceptible *S* (*DUX4* ^−^ D4T^−^), exposed *E* (*DUX4* ^+^ D4T^−^), infected *I* (*DUX4* ^+^ D4T^+^), resigned *R* (*DUX4* ^−^ D4T^+^), and dead *D*. Note the presence of 1 in the final row, showing that *D* (D4T^+^ myonuclear apoptosis) is an absorbing state, which a myonucleus cannot exit once it has entered. Having derived the transition matrix for the myonuclear model, we can calculate the mean absorption time *ω* (or in the context of our model, myonuclear life expectancy) from any given state via the fundamental matrix of the Markov chain [29, 30]–details in Appendix A. We are interested in how the model’s parameters describing *DUX4* expression affect FSHD disease progression from the starting “susceptible” or “healthy” state *S*, as this process incorporates all the parameters of *DUX4* mediated cytotoxicity (activation, transcription, translation, and downstream gene dynamics). The mean time to absorption in *D* when starting from state *S* is given by (derivation in Appendix A),

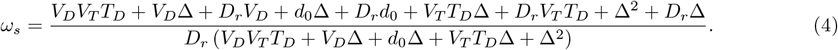

The quantity *ω*_*s*_ can be interpreted as the life expectancy of a myonucleus of an individual living with FSHD. Our framework allows us to express this lifespan in terms of the parameters underlying *DUX4* expression. Armed with this knowledge, we can investigate how changing each parameter of *DUX4* expression and function via therapy impacts myonuclear lifetime. This provides a way to prioritise anti-DUX4 therapy design by determining the most effective methods of prolonging FSHD-affected myonuclei survival within our model.

## III. RESULTS

### A. Limits of myonuclear lifetime

Equipped with the expression (4) we can study how the life expectancy of a myonucleus of an individual living with FSHD (*ω*_*s*_), depends on each parameter of the *DUX4* generation/function pathway. This provides insight into how a given anti-DUX4 therapy, which impacts some subset of the model parameters, may be expected to affect disease progression. The fold change in *ω*_*s*_ upon varying each parameter (*ω*_*s*_(*V*_*D*_), *ω*_*s*_(*d*_0_), …), while keeping all other parameters fixed at the levels recorded in Table I is presented in Figure 2**A** for a range of possible values. A large range of fold change in each parameter is investigated to show the limiting behaviour of changing each parameter (Figures 2**B**–**G**).

**Figure 2.**
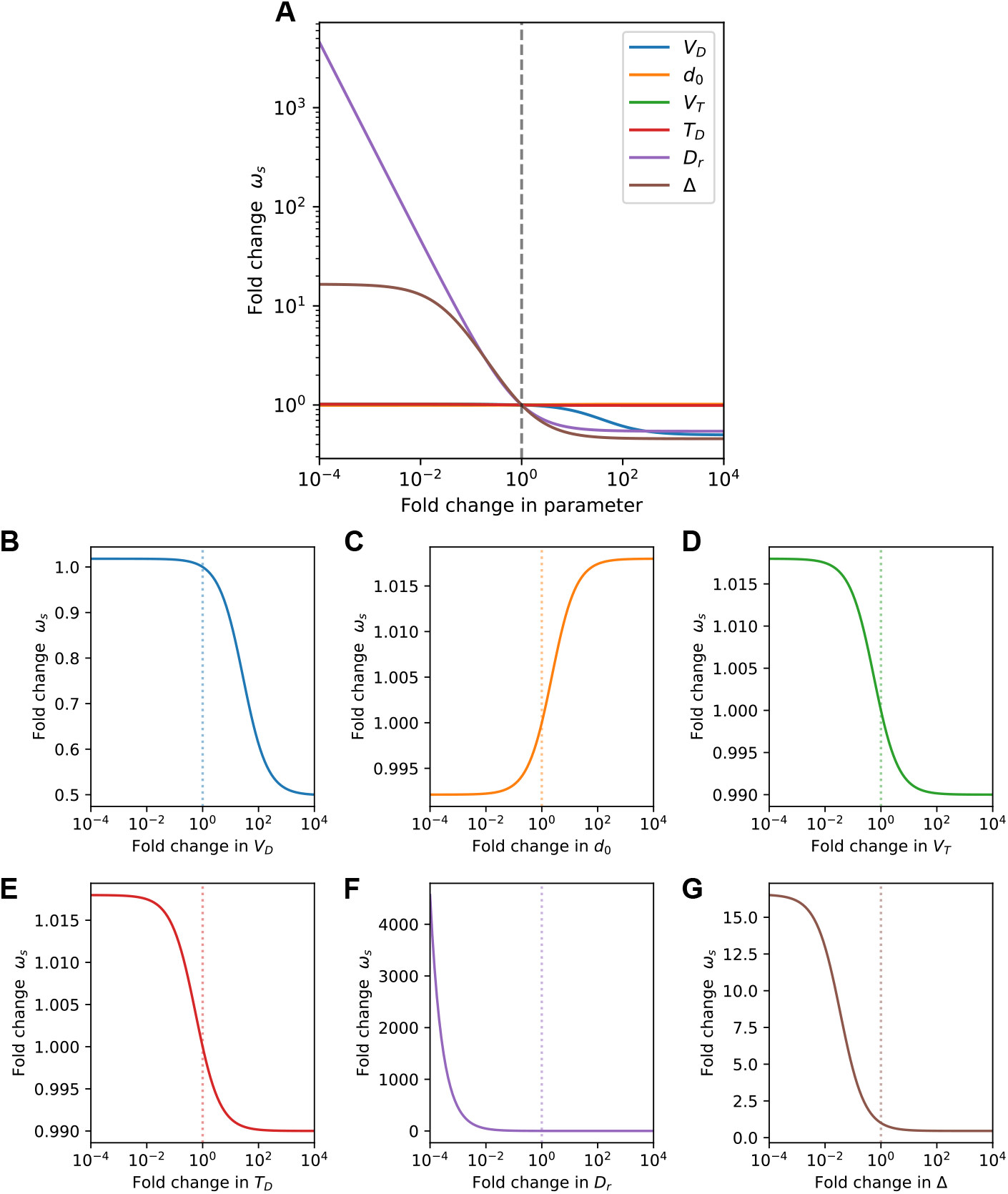
The lifetime of a myonucleus changes within the syncytial model of FSHD when parameters of the *DUX4* pathway are altered. (**A**) Fold change in the time to absorption in state *D* from state *S* (*ω*_*s*_) upon fold variation of the transcription rate of *DUX4* (*V*_*D*_), degradation rate of *DUX4* mRNA (*d*_0_), translation rate from *DUX4* mRNA to active DUX4 protein (*T*_*D*_), transcription rate of DUX4 target genes (D4Ts) (*V*_*T*_), death rate of D4T^+^ myonuclei (*D*_*r*_), and rate of DUX4 syncytial import (Δ). (**B**) to (**G**) show each rate parameter on individual axes, respectively. The dashed line on each plot indicates the endogenous case (no fold change in parameters). The vertical dashed lines in the centre of the plots indicate our estimates for the endogenous transition probabilities based on experimental data (Table I) [17].

All parameters except *D*_*r*_ (death rate of D4T^+^ myonuclei) have a sigmoidal relationship with *ω*_*s*_: the function is constrained by a horizontal pair of asymptotes and changing a parameter beyond threshold magnitudes has little impact on myonuclear life expectancy. The asymptotic values of *ω*_*s*_ under variation of each *DUX4* parameter, represent the absolute limit to which myonuclear life expectancy can be changed by varying that parameter by therapy. Equation (4) allows the derivation of these limits (Appendix B). For *V*_*D*_ we find:

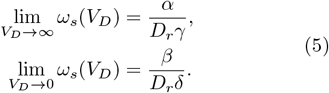

Where:

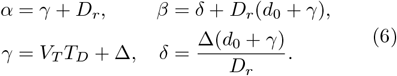

This demonstrates that to change the myonuclear life expectancy beyond these limits, *V*_*D*_ must be altered in tandem with other parameters as both the upper and lower bound of the myonuclear life expectancy with respect to *V*_*D*_ is dependent on other variables. The analytical form of these limits describes parameters that have positive or negative synergy with *V*_*D*_ with respect to *ω*_*s*_. For example, to increase the myonuclear life expectancy when *V*_*D*_ is very small 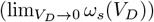, we can vary any of the other 5 variables with some degree of success. However, if *V*_*D*_ is very large 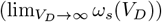, alterations in *d*_0_ will have no impact on the lower bound of the myonuclear life expectancy. This is understandable as *d*_0_ controls the backward rate of the transition between states *S* and *E*. If *V*_*D*_ = ∞, then increases in *d*_0_ will have no impact, as all myonuclei in state *S* will immediately be in state *E*. Biologically this is also intuitive as an extremely high transcription rate will generally mean that changes in a much smaller mRNA degradation rate will have minimal impact on reducing transcript abundance.

Taking limits helps identify why modifications of Δ (DUX4 syncytial import rate) result in such a large fold change in myonuclear life expectancy (Figure 2**G**). From Equation (4) it is found that (Appendix B),

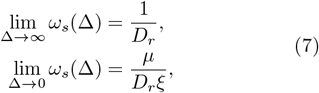

where:

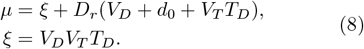

This shows that as the DUX4 syncytial import rate increases, the model becomes increasingly solely sensitive to *D*_*r*_. This is understandable as Δ effectively provides a “shortcut” in the *DUX4* pathway. Myonuclei can progress directly from state *S* to state *R*, if syncytial *DUX4* is imported into a nucleus, with zero dependence on any transcriptional or translational checkpoints. A situation with high Δ and *D*_*r*_ are therefore a ‘worst-case scenario” in the parameter space, with extremely rapid myonuclear apoptosis.

The other transcriptional and translational variables: *d*_0_, *V*_*T*_, *T*_*D*_ obey a similar relationship to *V*_*D*_ (Figures 2**C**–**E**) and have limiting functions dependent on a subset of the other parameters (Appendix B). The only parameter in the model that does not have a sigmoidal *ω*_*s*_ function is the DUX4 target gene-induced death rate *D*_*r*_. Taking limits (as shown in Appendix B), it is found that

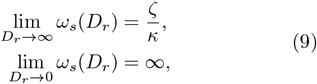

where:

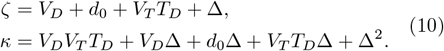

This confirms that *D*_*r*_ is the only parameter in the model with no upper bound to its impact on myonuclear lifetime. No matter how much we decrease *D*_*r*_, we would not find a plateau value of *ω*_*s*_, as encountered when varying another parameter (Figure 2**F**). This intuitively reflects that FSHD progression can effectively be immediately halted by completely blocking the apoptotic downstream consequences of *DUX4* and D4T positivity. No other parameter can achieve this effect as our model considers the therapeutic setting: we are studying perturbations of an ongoing disease state. If *V*_*D*_, for instance, is decreased to effectively zero, there is still some steady state reservoir of DUX4 in the system that can progress disease via the Δ term.

### B. Quantifying achievable efficacy

While the maximum achievable improvement in myonuclei survival has been quantified, of more relevance to drug discovery are the immediate gains that can be made by alterations of *DUX4* pathway parameters from their endogenous biological levels (Table 1). Figures 2**B**-**G** demonstrate that some parameters underlying *DUX4* expression, have endogenous values close to their asymptotic limits, meaning large changes in parameters are needed for small changes in myonuclear life expectancy, while for other parameters, this is not the case (the dashed lines in Figures 2**B**-**G**). Therefore the sensitivity of the myonuclear life expectancy to changes in each parameter from the endogenous level, alongside the maximal possible sensitivity has been quantified in Figure 3**A** (Appendix C). Interestingly, though *D*_*r*_ has the largest potential impact on the myonuclear lifetime, the immediate effect of altering Δ from the endogenous biological value is greater (Figure 3**B**). Combined with the fact that Δ is a less abstracted parameter than *D*_*r*_–many downstream effects of D4Ts are approximated into *D*_*r*_–this suggests that the most efficient variable of the *DUX4* pathway to target may be the DUX4 syncytial import rate. Turning to the other parameters, *V*_*D*_ is the next most efficient, with the rate of fold change of myonuclear lifetime around the endogenous biological value being close to the maximum for the parameter. With the *DUX4* pathway analysed on a parameter-by-parameter basis, we now consider that many therapies targeting the *DUX4* pathway will alter more than a single element, motivating studying interaction effects.

**Figure 3.**
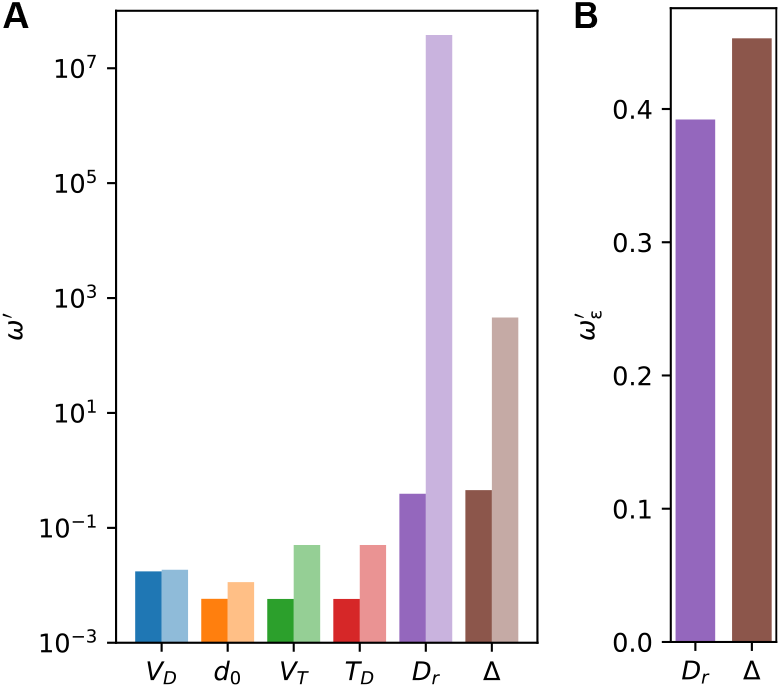
The impact of treatment depends on the endogenous behaviour of the system. (**A**) Sensitivity characterised by absolute fold derivatives of myonuclear lifetime (*ω′*: as calculated in Appendix C) with respect to each parameter in the endogenous case 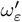 (dark bars) and the maximum of the function 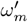 (light bars). (**B**) Linear plot of sensitivity of the endogenous myonuclear 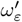 lifetime to Δ and *D*_*r*_.

### C. Interaction effects

An interaction effect is defined here as any combination of two given parameter values, which results in a greater-than-expected (super-additive) change in the myonuclear lifetime. For a given combination of parameters *i* and *j*, we denote the size of the interaction effect at a specific value of these two parameters by *I*(*x*_*i*_, *x*_*j*_). The matrix of interaction between all considered values of *i* and *j* is denoted **I**_*ij*_. The full details of the calculation of the term are given in Appendix D. The interaction effect can also be considered to be quantifying therapeutic synergy. The usefulness of characterising **I**_*ij*_ for a parameter combination is illustrated via Figure 4**A**, where there is a clear increase in the magnitude of *ω*_*s*_ when *V*_*D*_ and Δ are decreased in tandem. This is expected, as the plots in Figure 2**B** and **G** show the same behaviour when the parameters are varied singularly. However, what is not clear, is whether the increase in Figure 4**A** is above or below the expected linear combination of the parameters. ≈

**Figure 4.**
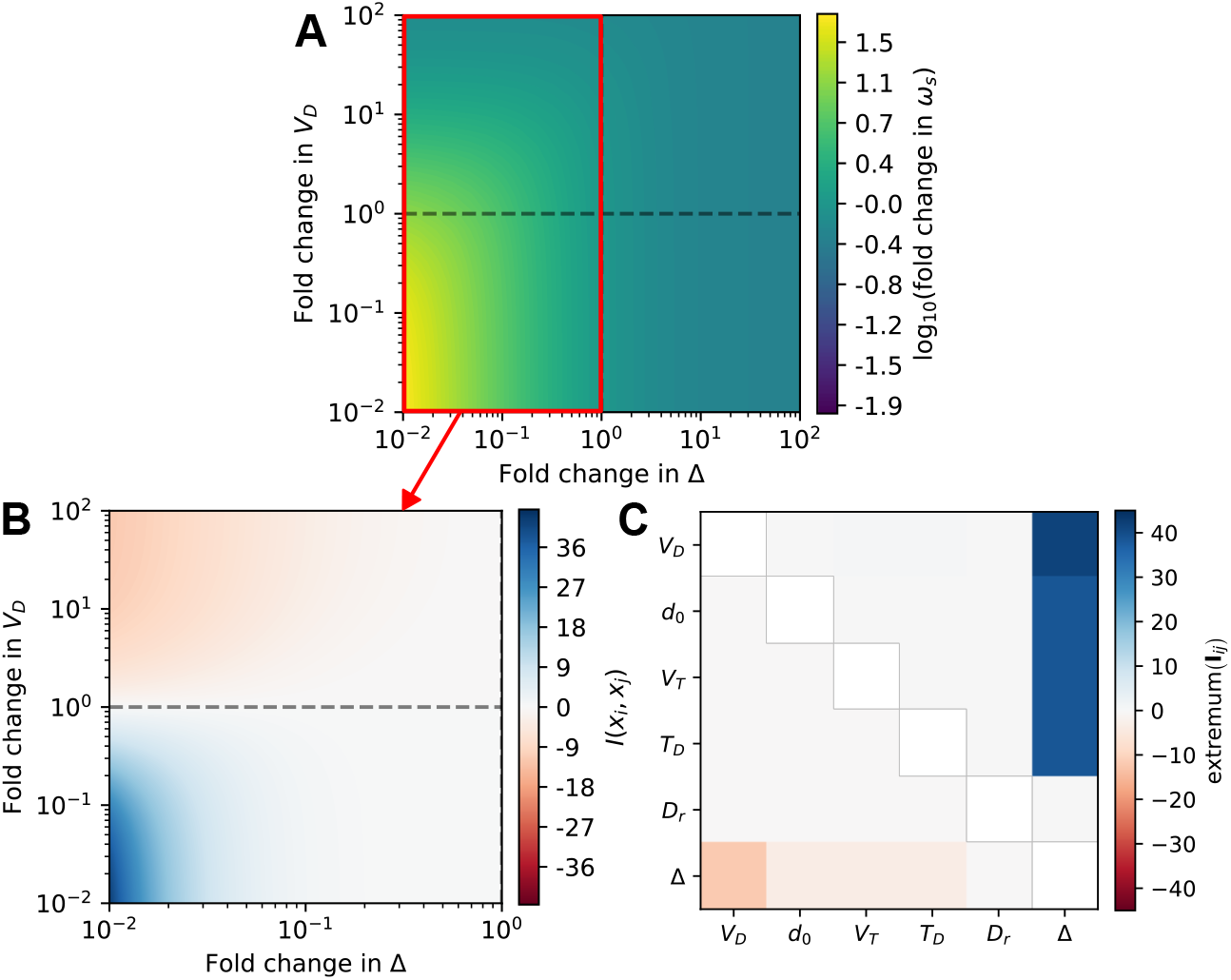
*DUX4* pathway parameters can have interaction effects. (**A**) Heat map showing the log10 fold change in the time to absorption in state *D* from state *S* (*ω*_*s*_) upon variation of the DUX4 syncytial import rate (Δ) and the *DUX4* transcription rate (*V*_*D*_). The red highlight indicates the area of interest plotted in Figure 4**B**. (**B**) Heat map showing the interaction effect of varying *V*_*D*_ with Δ as quantified via *I*(*x*_*i*_, *x*_*j*_) Equation (D1). (**C**) Heat map showing the extremum (either maximum or minimum) effect of each pairwise interaction **I**_*ij*_ (Equation (D2)) between *DUX4* pathway parameters in a 1 × 10^−2^–1 × 10^2^ range of fold change in each parameter. Below the diagonal colour indicates the minimum of **I**_*ij*_, above the diagonal colour indicates the maximum of **I**_*ij*_.

To examine this question, 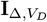 is plotted in Figure 4**B**. Immediately apparent is the strong interaction effect between the two parameters, both in a negative and positive fashion. Individually, reducing Δ or *V*_*D*_ by a factor of 10 returns an ≈13-fold increase or an ≈1.02-fold increase in the myonuclear lifetime, respectively. Together, reducing both Δ and *V*_*D*_ by a factor of 10 returns ≈55-fold increase. This dramatic synergy means there is an ≈42-fold additional additive increase in myonuclear lifetime obtained by applying both therapy strategies simultaneously.

To quantify the interaction effect across the entire considered parameter space, the minimum and maximum value of **I**_*ij*_ for each parameter combination is plotted in Figure 4**C**. The minimum and maximum are found via examining pairwise variations of parameter values over a range of 1 × 10^−2^–1 × 10^2^ fold changes in the endogenous value of each parameter. For most parameter combinations, there is a zero or near-zero interaction effect. Interestingly, only combinations of Δ with parameters other than *D*_*r*_ show synergy.

The reason for this can be seen in the schematic of the affected myonucleus in 1. In our model, the route which passes through the *E*–*I* transition and the route which uses the *S*–*R* transition are synergistic as they provide two separate pathways to reach myonuclear states susceptible to apoptosis. In the language of Markov chains, we have two closed loops which end in absorption. To cover the entire set of possible parameter combinations–including those where the number of parameters varied is *>*2–an interactive web application is available at: https://mcowley.shinyapps.io/fshdmcm/.

## IV. DISCUSSION

### A. Myonuclear coordination in FSHD

We have shown that in the context of our experimentally parameterised *in-silico* model, the rate of diffusion of DUX4 to other syncytial myonuclei is the most important parameter underlying *DUX4* expression for determining the time to myonuclear death. Myonuclear diffusion of DUX4 leading to a gradient of DUX4 and target gene expression has been shown in FSHD [31, 32]. Here we highlight that understanding the implications of such a mechanism is important to develop effective therapeutics.

From studies of syncytial dynamics, myonuclear coordination results in myonuclei becoming specialised. This can occur due to the location of the myonucleus in the syncitia–for example, being near a neuromuscular junction–or can be event-driven, such as after microleisons or severe injury [33]. From mechanistic studies of protein diffusion, the diffusion length– the mean distance a given protein travels before degradation–is controlled by a number of factors [34], and can be approximated via the distribution of the relative intensity of a fluorophore attached to the protein, with the rate of falloff of relative intensity away from the source myonucleus indicative of transport behaviour. Diffusion coefficients are key contributors to diffusion length and tend to be directly proportional to the molecular weight of the protein, with small proteins able to diffuse further than large proteins. The import coefficient for specific protein–myonuclei interactions is also important, as higher rates of nuclear import will result in a shorter diffusion length due to the protein being localised within nuclei, essentially acting as a protein sink. A counteracting mechanism to nuclear import is nuclear pore complex (NPC) diffusion; where proteins can diffuse out of the nucleus that has imported them. NPC diffusion appears to be of most relevance to small proteins, as the pore diameter restricts the size of protein that is able to pass through [34]. It has additionally been shown that myotube width influences diffusion length in a size-dependent manner [34]. Increasing the myotube width–without an accompanying increase in the myonuclear count– reduces the myonuclear density and causes the rate of import to effectively drop. This leads to extended diffusion for the large proteins unable to utilise NPC diffusion to escape the nuclear sinks. As highlighted by the mass dependence mechanisms of protein diffusion– and therefore myonuclear coordination–the expected behaviour of a system mediated by DUX4 diffusion will depend on its molecular properties.

DUX4 has a molecular mass of 45 kDa [35]. From the perspective of syncytial protein diffusion, this is a middling weight [34], making diffusion to, and import into, myonuclei 250 µm away possible. DUX4 would be expected to have low NPC diffusion due to its size– there is an exponential falloff in NPC diffusion rate with size [36]–making increases in myofibre circumference without a corresponding rise in myonuclear density effectively increase the diffusion length. This importance of the myonuclear density of the myofibre will therefore play an important role in determining the effect of protein diffusion in the *DUX4* pathway. Different muscle fibre types have different myonuclear densities depending on their myosin heavy-chain content, with slow-twitch fibres having higher densities and fast-twitch fibres having lower densities [37, 38]. This would make some fibre types more susceptible to the diffusion-mediated import of DUX4, as fast-twitch fibres with lower densities will have a longer diffusion length of DUX4. Therefore the impact of diffusion-mediated import of DUX4 will be greater in fast-twitch fibres. This idea aligns with findings that show enrichment of slow-twitch fibres in FSHD patients [39], and large fast-twitch restricted active force reductions in demembranated single muscle fibres from FSHD patients [40]. On the whole muscle scale, the influence of myonuclear coordination on disease progression would be related to that muscle’s fibre-type proportions. Assuming that the sporadic rate of DUX4 activation is the same for all myonuclei regardless of whether they are in fast-twitch or slow-twitch fibres, this mechanism could account for the heterogeneity of muscle pathology seen in FSHD.

### B. Interaction of *DUX4* expression events and diffusion

The other key result from our model is that there is a synergistic interaction effect between the rate of syncytial DUX4 diffusion and other *DUX4* expression parameters. Figure 5**A** shows a schematic diagram of a muscle fibre where some myonuclei are expressing D4Ts due to the primary expression of *DUX4*, denoted by the blue star and the fact that these myonuclei are *DUX4* ^+^ D4T^+^ (state *I*). Other myonuclei in the same fibre are *DUX4* ^−^ D4T^+^ (state *R*) due to the diffusion-mediated import of DUX4 having activated these genes without the myonucleus in question ever having expressed *DUX4* itself.

**Figure 5.**
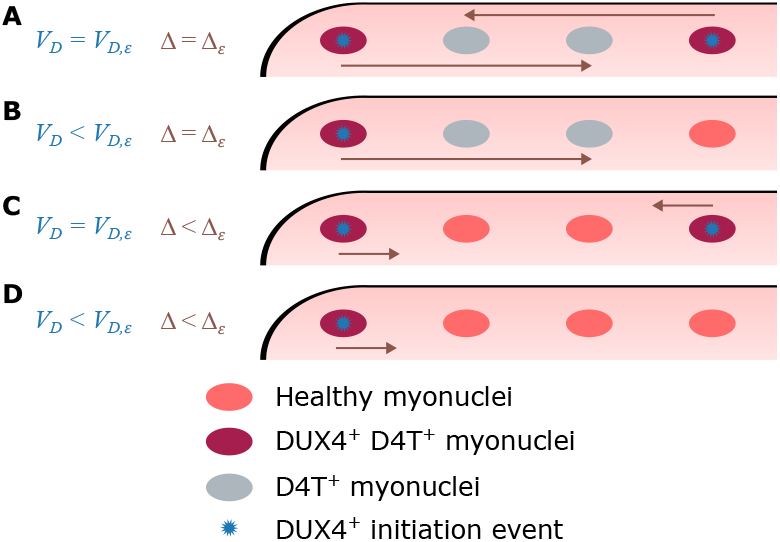
Perturbation of the rate of *DUX4* transcriptional events and the rate of the diffusion-mediated import of DUX4 has synergistic effects. (**A**) The *DUX4* transcription rate *V*_*D*_ and DUX4 syncytial import rate Δ are at their endogenous levels (*V*_*D,ε*_ and Δ_*ε*_). *DUX4* expression events (blue stars) occur across the myonuclei in the myofibre and diffusing DUX4 (brown arrow) activates DUX4 target genes (D4Ts) in neighbouring myonuclei. (**B**) Reducing *V*_*D*_ from the endogenous level *V*_*D,ε*_ results in fewer *DUX4* expression events (blue stars) across the myonuclei in the myofibre. Additional D4T^+^ myonuclei are still created via diffusion-mediated import. (**C**) Reducing Δ from the endogenous level Δ_*ε*_. *DUX4* expression events occur randomly across the myofibre, but no diffusion-mediated import and D4T activation in neighbouring myonuclei occurs. (**D**) Both *V*_*D*_ and Δ are reduced. There are fewer *DUX4* expression events and an initiating positive myonucleus does not affect nearby myonuclei.

As an example treatment scenario (Figure 5**B**), perturbation of the muscle fibre has reduced the magnitude of *V*_*D*_ such that only a single myonucleus undergoes a *DUX4* ^+^ primary expression “initiation event”.

However, despite this event only occurring once, a large number of neighbouring myonuclei in the fibre acquire the D4T^+^ transcriptional state due to diffusion-mediated import of DUX4 (brown arrow). If instead, a perturbation alters the magnitude of Δ (Figure 5**C**), the influence of a single *DUX4* ^+^ myonucleus is smaller, as the change in diffusion length reduces the number of myonuclei that import the pathological protein. In this scenario, *V*_*D*_ is still at the endogenous level, and due to this randomly distributed myonuclei become *DUX4* ^+^ D4T^+^ with no spatial relationship. Both *V*_*D*_ and Δ being reduced (Figure 5**D**) creates the ideal scenario where *DUX4* ^+^ expression events are both rare and have minimal impact on the surrounding myonuclei. The *DUX4* ^+^ expression event route and the diffusion-mediated import route can be considered separate sub-pathways within the overall DUX4-mediated dysregulation chain.

Interestingly, due to the clear difference in *DUX4* ^+^ D4T^+^ arrangement in the two settings, spatial single-nuclear transcriptomic experiments can answer the question of which of these two pathways is the more important. If there are myonuclei in state *I* and *R* randomly distributed across the myofibre, likely the primary expression of *DUX4* is the dominant factor. Whereas, if there are predominantly clusters of *I* and *R* myonuclei, the diffusion-mediated import route has a greater impact. Recent studies suggest that clusters of D4T^+^ myonuclei can form in myotubes, with enrichment of FSHD phenotypic myonuclei in myotubes over unfused myocytes [41]. By inspecting the observed distributions in a probabilistic manner, it will be possible to validate the predictions made in this work and infer the parameters of the *DUX4* -pathway in other developmental states and genotypic settings.

## V. SUMMARY

Mathematical models of biological systems provide the opportunity to perturb the system explicitly. Applying this framework to analytical solutions of statistical models allows distributional inferences about stochastic events. Here, we have taken this approach with a Markov chain model of FSHD and found that protein diffusion may be a parameter to which myonuclear lifetime in the FSHD setting is very sensitive. Surprisingly, when altered by treatment in a biologically relevant parameter regime, syncytial diffusion is found to be more impactful than mediating D4T apoptosis-inducing effects. Protein diffusion may even account for the heterogeneity of muscle pathology, with varied myofibre-type compositions resulting in disparate outcomes.

Additionally, we found that there is a synergistic interaction effect between DUX4 diffusion and parameters involved in *DUX4* and D4T transcription/translation. This suggests that therapeutic strategies that consider both *DUX4* transcriptional events and the rate of diffusion-mediated import of DUX4 will be most effective. A testable hypothesis for future spatial single-nuclear transcriptomic studies is provided in the form of a prediction of two possible states of the spatial distribution of *DUX4* ^+^ and/or D4T^+^ myonuclei, and a computational tool to screen combinatorial therapeutic treatments is made freely available.

## ACKNOWLEDGMENTS

MVC was supported by Friends of FSH Research (Project: An *in-silico* approach to understanding *DUX4* expression). CSRB was supported by the Turing-Roche Partnership. The Zammit lab was also generously supported by the Medical Research Council (MR/S002472/1), SOLVE FSHD, FSHD Society and Association Francaise contre les Myopathies.

## SOFTWARE AVAILABILITY

A web app to exhaustively test parameter combinations for pre-clinical screening is provided here: https://mcowley.shinyapps.io/fshdmcm. Instructions to self-host the Shiny-based web app and recreate the plots in Figures 2–4 can be found here: https://github.com/MVCowley/fshdmcm.

## Appendix A: Calculating mean absorption times

For a given probability matrix in an absorbing Markov chain, there is a sub-matrix *Q* containing only the *t* transient states [29, 30]. We have 4: *S, E, I*, and *R*. From *Q* it is possible to find the fundamental matrix *N* via,

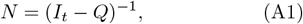

where *I*_*t*_ is the 4 × 4 identity matrix. Performing the matrix inversion via Gauss-Jordan elimination it is found that,

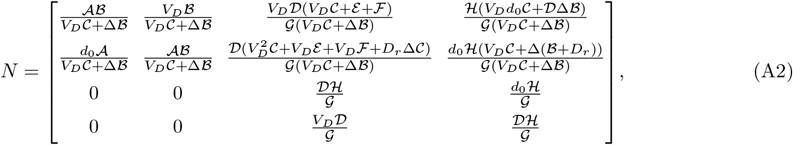

where for readability, the following substitutions have been made:

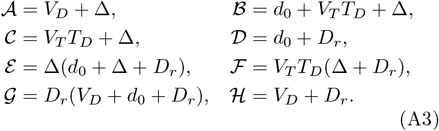

With the fundamental matrix in hand, the mean time to absorption *ω*_*l*_ in state *D* from each transient state *l* can be found via,

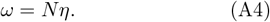

Where *ω* is a vector of all absorption times of length *t* and *η* is a length *t* column vector of the expected waiting times (*η*_*l*_, …, *η*_*t*_) in each state as defined by the exponential distribution,

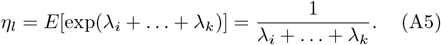

Thus *ω*_*s*_ in Equation (4) is given by the first element of the vector *ω*.

## Appendix B: Mean absorption times and limits with respect to selected parameters

### 1. *DUX4* transcription rate

With some rearrangement with respect to *V*_*D*_, Equation (4) can be rewritten as,

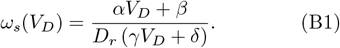

Where the other parameters have been subsumed into the constants,

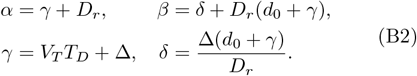

The derivation of limits for the relationship of *V*_*D*_ with *ω*_*s*_ is shown in § III A.

### 2. DUX4 syncytial import rate

With some rearrangement with respect to Δ, Equation (4) can be rewritten as,

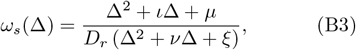

where,

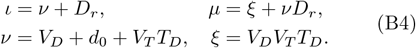

The derivation of limits for the relationship of Δ with *ω*_*s*_ is shown in § III A.

### 3. DUX4 target gene positive myonuclear apoptosis rate

With some rearrangement with respect to *D*_*r*_, Equation (4) can be rewritten as,

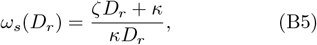

where,

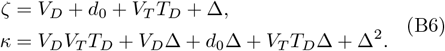

The derivation of limits for the relationship of *D*_*r*_ with *ω*_*s*_ is shown in § III A.

### 4. *DUX4* mRNA degredation rate

With some rearrangement with respect to *d*_0_, Equation (4) can be rewritten as,

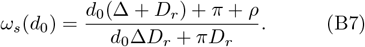

Where the other parameters have been subsumed into the constants,

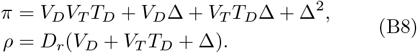

Limits may then be taken at zero and ∞, finding that,

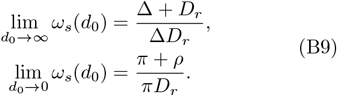

Therefore, *d*_0_ has a sigmoidal relationship with *ω*_*s*_.

### 5. DUX4 target transcription rate and *DUX4* mRNA translation rate

With some rearrangement with respect to *V*_*T*_, Equation (4) can be rewritten as,

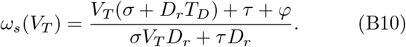

Where the other parameters have been subsumed into the constants,

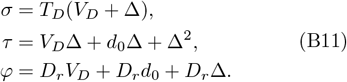

Limits may then be taken at zero and ∞, finding that,

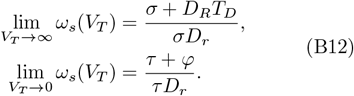

By substituting all occurrences of *V*_*T*_ in the above equations for *T*_*D*_, and all occurrences of *T*_*D*_ for *V*_*T*_, an identical set of equations and limits can be found for *ω*_*s*_(*T*_*D*_). Therefore, both *V*_*T*_ and *T*_*D*_ have a sigmoidal relationship with *ω*_*s*_.

## Appendix C: Quantification of parameter sensitivity

To understand how small changes in model parameters alter the myonuclear lifetime (*ω*_*s*_), a metric *ω′*– which uses the gradient of the relationship between the parameter of interest and *ω*_*s*_–is established.

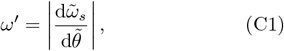

Where *θ* is a parameter in the set of Markov chain parameters and 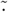 is a quantity normalised by its endogenous value presented in Table I e.g. 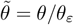. To simplify notation, *ω′* uniquely refers to the absolute fold derivative of the myonuclear lifetime with respect to the starting state *S*, as opposed to a vector of the quantity from each state. Instead, the subscript is used to denote the point in the *ω*_*s*_–parameter relationship curve: 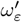 for the endogenous value (at the dashed line in Figure 2) and 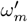 for the maximum value 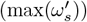).

Only the range of parameters considered in Figure 2 is used to determine the maximum, as this avoids the value of 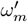 for *D*_*r*_ being infinity and captures the region of sigmoidal behaviour for the other parameters. The absolute value of the derivative is used to allow easier comparison between *d*_0_, which has an increasing monotonic relationship with myonuclear lifetime, and the other parameters which have a decreasing monotonic relationship. The endogenous case, 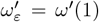, is important as it represents the initial impact of changing a given rate from its biologically estimated level.

## Appendix D: Quantifying interaction

An interaction effect is defined in this work as any combination of two given parameter values, which results in a greater than expected (super-additive) change in the myonuclear lifetime–where the expected change is the sum of the calculated myonuclear lifetimes when varying the given parameters independently, offset such that in the endogenous case *ω*_*s*_(*θ*_*ε*_) the interaction is normalised to zero. This can be achieved for any given point in the pairwise parameter space via,

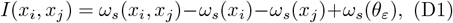

where *x*_*i*_ and *x*_*j*_ are the values of the parameters *i* and *j* the interaction term is being computed for, and *θ*_*ε*_ is the endogenous parameter set. Hence, when {*x*_*i*_, *x*_*j*_} ⊂ *θ*_*ε*_, *I*(*x*_*i*_, *x*_*j*_) = 0.

The interaction term is computed as a fold change to make the values obtained comparable to the plots in Figure 2. For each parameter combination a matrix of interaction terms **I**_*ij*_ is computed by,

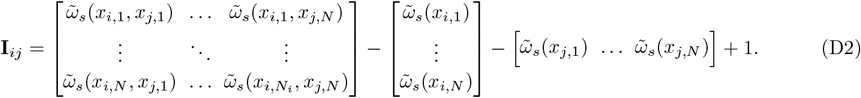

Where 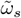 is *ω*_*s*_ normalised by *ω*_*s*_(*θ*_*ε*_), and *N* is the number of elements in the range of parameter values for *i* and *j*. Thus each index of **I**_*ij*_ is the additional additive fold change of altering two parameters together over the expected fold change of adjusting the parameters.

